# Cell-specific variant-to-gene mapping identifies conserved neural and glial regulators of sleep

**DOI:** 10.64898/2026.04.07.715910

**Authors:** A.J. Zimmerman, S. Biglari, K.B. Trang, E. Almeraya Del Valle, A.I. Pack, S.F.A. Grant, A.C. Keene

**Affiliations:** Department of Genetics, University of Pennsylvania Perelman School of Medicine, Philadelphia, PA, 19104, USA; Department of Medicine, University of Pennsylvania Perelman School of Medicine, Philadelphia, PA, 19104, USA; Center for Spatial and Functional Genomics, Children’s Hospital of Philadelphia. Philadelphia. PA, 19104, USA; Department of Pediatrics, University of Pennsylvania Perelman School of Medicine, Philadelphia, PA, 19104, USA; Divisions of Human Genetics and Endocrinology & Diabetes, Children’s Hospital of Philadelphia. Philadelphia. PA, 19104, USA; Department of Biology, Texas A&M University, College Station, TX, USA

## Abstract

Excessive daytime sleepiness (EDS) is a heterogeneous phenotype with little known of its genetic basis. Large-scale genome-wide association studies (GWAS) have reported genomic loci associated with EDS, though since most of these are non-coding, the causal gene(s) underlying the association are not known. Additionally, the cell types in which these genes exert their effects on sleep have not been functionally explored *in vivo*. Here, we employed a chromatin-based variant-to-gene mapping approach to first implicate candidate effector genes at EDS GWAS loci in human-derived neural and glial cell lines. Subsequent cell type-specific RNAi knockdown of orthologous genes using neural and glial GAL4 drivers in *Drosophila* confirmed cell-specific regulation of sleep by these GWAS-implicated effector genes. Among these, *ruby* (ortholog to *AP3B2*), a component of the AP-3 vesicular trafficking complex emerged as a robust sleep regulator. Targeted knockdown in flies localized *ruby* function to astrocyte-like glia, where loss of *ruby* increased sleep duration. The conserved role of *ruby/ ap3b2* was validated in zebrafish where CRISPR-mediated loss increased daytime sleep. Together, these findings show that physical variant-to-gene mapping predicted cell-type–specific gene function for complex sleep traits and revealed *ruby/AP3B2* as a conserved glial regulator of sleep and arousal. This work provides a generalizable framework for connecting non-coding GWAS variants and their corresponding effector genes to identify novel and highly conserved regulators of sleep.

## Introduction

Dysregulation of sleep duration, timing, and quality is associated with increased risk for negative health outcomes including cardiometabolic disease and mortality [1,2]. Given a strong heritable component [3], there is an unmet need to identify the causal genetic and molecular drivers of poor sleep to improve overall health. Significant efforts have been made to define the underlying genetic drivers of sleep disorders, including insomnia, sleep apnea, and excessive daytime sleepiness (EDS) through genome wide association studies (GWAS) in large population cohorts[3–10]. However, few candidate genes identified through GWAS have been functionally validated, and the genetic and cellular bases of sleep disorders remain poorly understood.

While the field of sleep genetics has primarily focused on insomnia, EDS is also associated with negative health outcomes including poor metabolic and cognitive health [11,12]. Relative to other sleep disorders, EDS has received little attention given it is widely considered a heterogeneous symptom rather than a single disorder. Previous GWAS efforts showed that defined broadly EDS can be genetically subdivided into two biologically distinct subtypes, sleep propensity and sleep fragmentation [3]. While the genetic associations for each of these biological subtypes of EDS point to distinct underlying mechanisms, these observations have not been functionally validated. Mechanistic characterization of these loci should reveal causal effector genes contributing to EDS and in turn improve our understanding of the etiology.

Many GWAS efforts have reported genetic loci associated with sleep regulation. However, defining the underlying causal effector genes at these loci remains a central challenge for the field [13].Given the complex genetic architecture regulating sleep and its polygenic nature [14], mechanistic testing of multiple candidate genes in animal models can be costly and challenging. Further complicating interpretation, more than 90% of GWAS loci reside in non-coding regions of the genome [13,15]. Consequently, genes selected for functional validation using the common method of nearest-gene assignment often do not correspond to the true causal effector genes tagged by GWAS [16,17].

Because the majority of GWAS signals reside in non-coding regions, many variants localize to cis-regulatory elements, including enhancers and promoters, that shape spatiotemporal gene expression programs [15,18–21]. Importantly, the activity of these regulatory elements is often highly cell-type specific, underscoring the need to interpret GWAS variants within relevant cellular contexts[13,22]. Cell-specific chromatin-based variant-to-gene mapping provides a powerful framework for defining how regulatory variants influence gene function across distinct cell types, particularly for variants that are not broadly active across tissues [20,23–26]. Here, we apply chromatin capture techniques across five neural and glial cell types to identify three-dimensional genomic interactions between GWAS-implicated putative causal variants and their candidate effector genes, thereby nominating gene regulatory interactions that contribute to the heritability of EDS.

To functionally validate genes implicated in this variant-to-gene mapping from EDS GWAS, we employed a cross-species screening approach. Non-mammalian genetic model organisms have provided critical insight into the genetic regulation of sleep [27,28]. Both *Drosophila melanogaster* (fruit fly) and *Danio rerio* (zebrafish) exhibit the canonical behavioral hallmarks of sleep, and genetic perturbation studies in these systems demonstrate that core sleep-regulatory mechanisms are highly conserved with those of mammals [27,28]. While neuronal genes have been the principal focus of sleep regulation, there is growing evidence that multiple glial cell types also play critical and conserved roles in sleep and circadian control [29–35]. Consequently, mechanistic insight into the coordinated contributions of sleep-regulating neurons and glia is essential for understanding the etiology of human sleep disorders. Informed by our cell-specific variant-to-gene mapping, we performed cell-type-specific RNAi knockdown in *Drosophila*, revealing unique neural and glial roles for each human GWAS-implicated gene. We ultimately validated our findings in zebrafish providing a thorough cross-species characterization for a novel gene not previously implicated using nearest-gene approaches, demonstrating the capacity for gene discovery using this approach.

## Results

### 3D genomic mapping links sleep-associated variants to distal target genes

To implicate effector genes at EDS GWAS loci, we leveraged sentinel signals from a previous GWAS report for self-reported daytime sleepiness [3] (**Fig. 1A**). This previous GWAS effort clustered variants into two biological subtypes of EDS using accelerometry data, one marked by high sleep propensity (e.g. hypersomnia) and one defined by sleep fragmentation. Given EDS is a consequence of poor sleep quality and is associated with significant morbidity, we elected to focus our variant-to-gene efforts on this biological subtype, effectively uncoupling this trait from the distinct genetic architecture of intrinsic hypersomnia.

**Figure 1.**
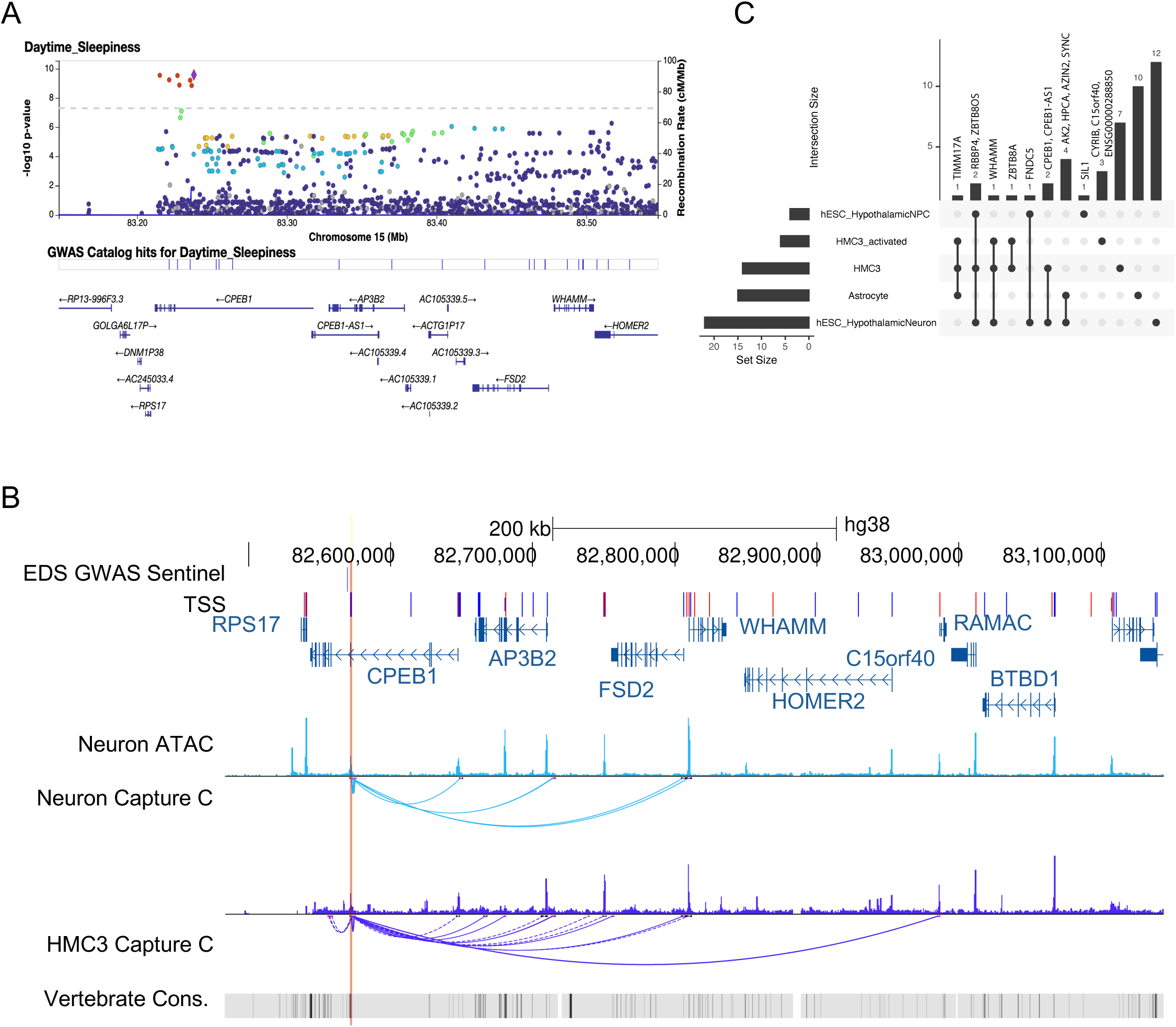
Variant-to-gene mapping nominates cell-specific effector genes for daytime sleepiness. **A.** Locus zoom plot of the GWAS association on chromosome 15 for excessive daytime sleepiness (from https://sleep.hugeamp.org/). Sentinel variant is shown as purple diamond and proxy variants are colored by linkage disequilibrium *r^2^* value. **B.** Variant-to-gene mapping at the chromosome 15 locus in iPSC-derived hypothalamic neurons and HMC3 microglia. EDS GWAS sentinel (lead) variant is displayed and proxy variant is indicated by red highlight. Proxy variant resides in an intron of *CPEB1* and forms chromatin contacts (capture C) with accessible (ATAC) promoters (TSS) of multiple distal genes that differ by cell type. Conservation is shown (vertebrate cons.) **C.** Upset plot showing number of target effector genes identified in each cell type.

From the 27 sentinel signals associated with the EDS-sleep fragmentation cluster [3], we performed LD expansion for proxy variants (*r^2^*>0.8, details in Methods), then applied variant-to-gene mapping by intersecting variants with cell-type specific cis-regulatory elements (cREs) – defined as genomic regions exhibiting open chromatin (ATAC-seq peaks) and simultaneously contacted by a chromatin interaction (Hi-C or Capture-C loops) as previously described [23] (**Fig. 1B, Supplementary Fig. 1, and Supplementary Table 1**). We performed this analysis across five different cell types including embryonic stem cell-derived hypothalamic neural progenitors and mature hypothalamic neurons, resting and activated HMC3 microglia, and astrocytes. This approach yielded 45 unique candidate effector gene promoters (33 of which are protein-coding) across all 5 cell types contacted by 35 proxies residing in cREs resulting from 10 lead sentinel signals (**Figure S1 and Supplementary Table 1**). Mature hypothalamic neurons yielded the most unique candidate effector genes, with 12 promoter-proxy SNP interactions, followed by 10 in astrocytes, 7 in resting HMC3s, 3 in activated HMC3s, and 1 in hypothalamic neural progenitors (**Fig. 1C**).

Pan cell type analysis showed sharing of at least one effector gene (**Fig. 1C**). Of the 10 sentinel signals, it was noted that the rs2787120-tagged locus and its proxies contacted a dense cluster of 12 genes on chromosome 1p35.1 (*AK2, S100PBP, TMEM54, HPCA, YARS1, AZIN2, BSDC1, ZBTB8A, FNDC5, SYNC, RBBP4, and ZBTB8OS*) within open chromatin regions in both neural and glial types (**Figure S1 and Supplementary Table 1**). Despite only five high-LD proxies for the rs17356118-tagged locus, a diverse cluster of 14 genes on chromosome 15q25.2 was implicated that includes key synaptic modulators (*CPEB1*, *HOMER2*, *AP3B2*) and multiple antisense/non-coding RNAs (*CPEB1-AS1*, *TMC3-AS1*, *SNHG21*) (**Fig. 1B**, **Figure S1 and Supplementary Table 1**). Across all five cell types, we implicated 22 genes with glial-specific contacts, 15 genes with neural-specific contacts and 8 genes with both glial and neural contacts.

### Cell-specific *Drosophila* screen of candidate effector genes

To test candidate genes identified through our variant-to-gene approach for their contribution to sleep regulation, we performed cell-type-specific knockdown in *Drosophila* of these genes in either neurons or glia, guided by variant-to-gene mapping that predicts the relevant cellular context of action.

We first identified orthologous genes using DIOPT (**Figure S2**). Of the 45 candidate effector genes, 21 had orthologs in *Drosophila*. Of these, 13 RNAi lines were screened using pan-neuronal (*nSyb*) and 14 RNAi lines were screened using pan-glial (*repo*) GAL4-UAS (**Fig. 2A**). This resulted in the identification of multiple genes that suppressed or increased sleep in each cell type. Notably, pan-neuronal knockdown of the ortholog to *HPCA*, *neurocalcinin* (*nca*), suppressed sleep (**Fig. 2B**), confirming previous reports that this gene is sleep-promoting [36]. Similarly, pan-neuronal knockdown of *orb* (ortholog to *CPEB1*) led to short sleep, while knockdown of *TyrRs* (*YARS1*), *idit* (*FNDC5*), *sil1* (*SIL1*), *odc2* (*AZIN2*), *archease* (*ZBTB8OS*), *CG4537* (*CRIPT*), and *CG8916* (*GABRA4/GABRB1*) increased sleep compared to both nSyb-GAL4>UAS-Luciferase and nSyb-GAL4/+ controls (**Fig. 2B**). In glia, knockdown of genes associated with the chromosome 15q25.2 locus, *CG14966* (C15orf40), *homer* (*HOMER2*), and *ruby* (*AP3B2*) increased sleep (**Fig. 2C**). Additionally, glial knockdown of two zinc finger and BTB domain containing genes at the 1p35.1 locus, *archease (ZBTB8OS) and promyleocytotic zinc finger* (plzf/ *ZBTB8A*) also increased sleep compared to both repo-GAL4>UAS-Luciferase and repo-GAL4/+ controls (**Fig. 2C**). The relatively high hit rate compared to previous unbiased sleep screens suggests variant-to-gene mapping based on sleep GWAS enriched for sleep-regulating genes in *Drosophila* [37–39]. It should be noted that orthologs to multiple genes implicated by the human chromosome regions 1p35.1 and 15q25.2 altered sleep, suggesting multiple causal genes at these loci.

**Figure 2.**
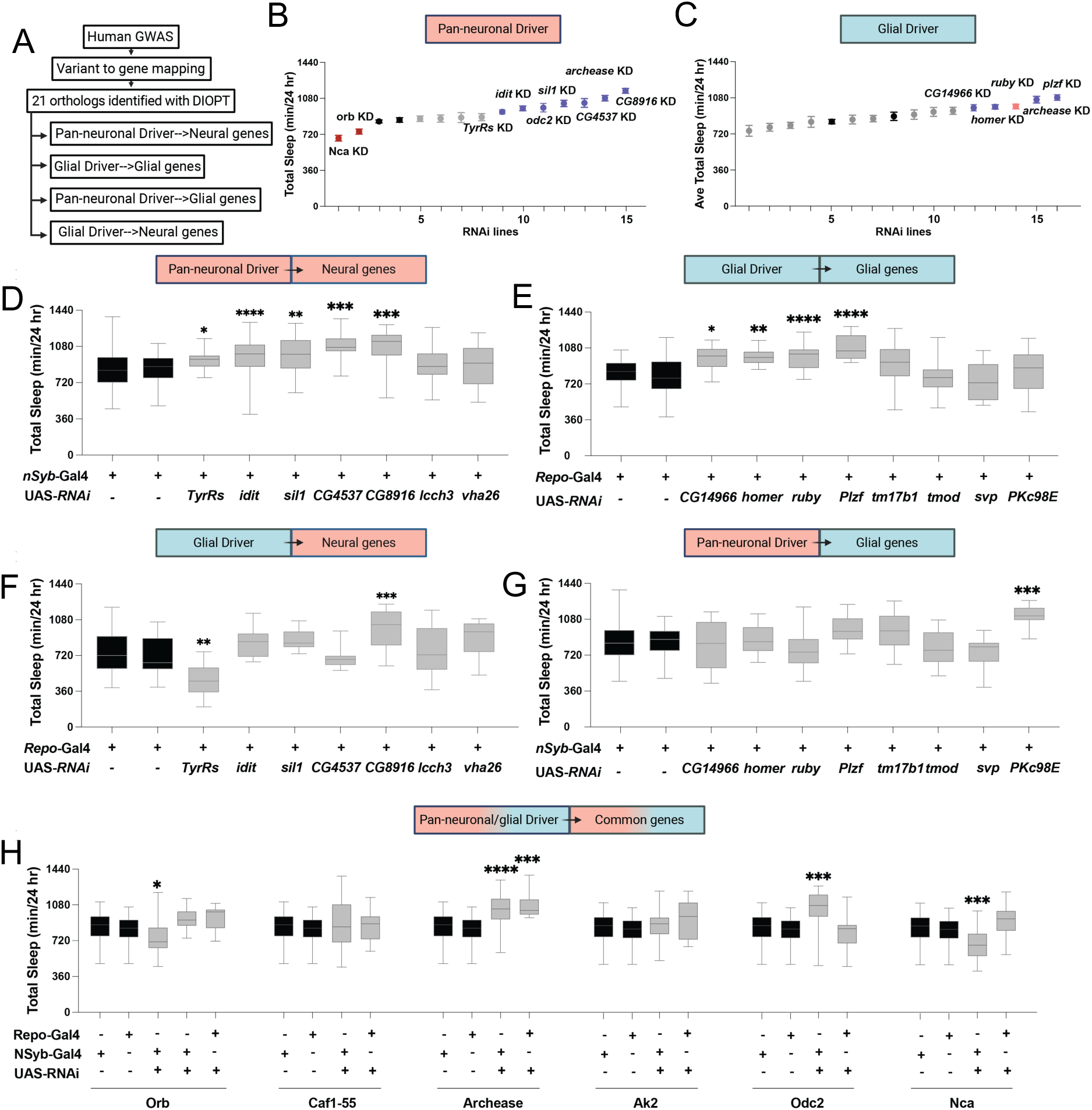
RNAi screen in *Drosophila* defines cell-specific regulation of excessive sleepiness. **A.** Schematic of orthologous-based RNAi screen. Candidate genes predicted from human GWAS variant-to-gene mapping were identified using DIOPT and tested in *Drosophila* using RNAi expressed in neurons (*nSyb*-GAL4) or glia (*repo*-GAL4). **B.** Total sleep minutes over a 24-hour period in viable RNAi crosses expressed selectively in neurons under control of *nSyb*-GAL4 (13 lines, n>16 per line). Black dots indicate control sleep responses, while colored dots highlight selected RNAi lines. Knockdown of *nca* and *orb* produced short sleep phenotypes, whereas knockdown of *TyrRs*, *idit*, *sil1*, *odc2*, *archease*, *CG4537*, and *CG8916* increased sleep compared to both *nSyb*-GAL4 > UAS-luciferase and *nSyb*-GAL4/+ controls. **C.** Total sleep minutes over a 24-hours period in viable RNAi crosses expressed selectively in glia under the control of *repo*-GAL4 (14 lines, n>16 per line). Knockdown of the *CG14966*, *homer*, *ruby*, *archease*, and *plzf* increased sleep compared to both *repo*-GAL4 > UAS-*luciferase* and *repo*-GAL4/+ controls. **D.** Pan-neuronal knockdown of the candidate genes predicted to function in neurons significantly altered sleep duration in comparison to controls (one-way ANOVA: F_8,357_= 9.210, p < 0.0001). The pan-neuronal knockdown in *TyrRs* (p < 0.05), *idit* (p < 0.001), *sil1* (p < 0.001), *CG4537* (p < 0.001), and *CG8916* (p < 0.0001) significantly increases sleep relative to controls. **E.** Glial knockdown of the candidate genes predicted to be function in glial cells significantly affects sleep duration (one-way ANOVA: F_9,243_= 9.624, p < 0.0001). The glial knockdown in *CG14966* (p < 0.01), *homer* (p < 0.001), *rb* (p < 0.0001), and *plzf* (p < 0.001) significantly increases sleep compared to controls (black boxes). **F.** Effects of knocking down neuronal genes in glia on sleep duration was less robust (ANOVA: F_9,174_= 7.316, p < 0.0001) with only one line with increased sleep *CG8916* (p < 0.0001), and one line with decreased sleep *TyRs* (p < 0.001) relative to controls. **G.** Effects of knocking down glial genes in neurons was less robust on sleep duration (ANOVA: F_10,335_= 6.065, p < 0.0001) with only one line with increased sleep *Pkc98E* (p < 0.0001) relative to controls. **H.** Genes predicted to function in both neurons and glia differentially regulate sleep depending on the driver used (ANOVA: F_18,621_= 7.730, p < 0.0001). The pan-neuronal and glial knockdown in *orb* (p < 0.01), *Archease* (p < 0.0001), *Odc2* (p < 0.0001), and *Nca* (P <0.0001) significantly modulate sleep. Lines and error bars represent mean ± SEM; Boxes represent median and quartiles. Statistical comparisons were performed relative to both GAL4 and UAS controls. * p < 0.05, ** p < 0.01, *** p < 0.001, **** p < 0.0001 represent adjusted p-value following Dunnett’s multiple comparisons test.

Several genes identified through our variant-to-gene mapping approach were specifically implicated in neural cells only or glial cells only, while others were common to both. To determine the cell-specific roles of these genes, we examined sleep phenotypes resulting from neural-specific knockdown of neural genes followed by glial-specific knockdown of glial genes. For genes specifically implicated through variant-to-gene mapping in neurons only, *nSyb*-GAL4>RNAi resulted in increased sleep in 5/7 lines tested (**Fig. 2D and Figure S2**). For genes specifically implicated as operating in glia only, *repo*>RNAi resulted in increased sleep in 4/8 lines (**Fig. 2E and Figure S2**). Thus, when the human cell type used for identification was matched to the *Drosophila* driver (e.g. neural – *nSyb* or glial – *repo*) the resulting phenotype was excessive sleep at least 50% of the time.

To validate cell-type specificity, we next tested the effects of knocking down neuron-specific drivers in glia and vice versa. When the GAL4 driver was discordant to the cell type in which the effector gene was implicated, the effects were less robust. When neural genes were knocked down with *repo*-GAL4, 1/7 increased sleep, and 1/7 decreased sleep (**Fig. 2F and Figure S2**). When glial genes were knocked down with *nSyb*-GAL4, 1/8 increased sleep (**Fig. 2G and Figure S2**). We next tested the seven genes that were common to both neurons and glia using both drivers. For the genes *orb* (*CPEB1*) *and nca* (*HPCA*), knockdown in glia yielded an opposite, but non-significant, phenotype to neuronal knockdown (**Fig. 2H and Figure S2**), suggesting that these genes influence sleep in a cell-specific manner. Only with *archease* (*ZBTB8OS*) was sleep significantly increased with both neuronal and glial knockdown (**Fig. 2H and Figure S2**). These results indicate sleep regulators have unique functions in neurons and glia and support the hypothesis that cell type-specific variant-to-gene mapping enriches for functional regulation of sleep.

We next sought to further examine the role of *ruby* (AP3B2) in sleep regulation given the robustness of the sleep-promoting phenotype resulting from knockdown and the potential functional novelty of its molecular role in sleep. *AP3B2* was also only implicated in glia and selectively affected sleep with glial knockdown. *ruby* encodes the beta3-adaptin subunit of the Adaptor Protein-3 (AP-3) complex that has been implicated in lysosome, synaptic vesicle, and protein trafficking; yet this complex has not been previously implicated in sleep regulation [40–42]. To validate findings from the initial screen and confirm they were not due to off-targets effects, we repeated pan-glial knockdown with multiple independent RNAi lines. For both lines tested, *ruby* knockdown flies consistently slept more than controls harboring the *repo*-GAL4 or UAS-RNAi lines alone (**Fig. 3A-C**). We quantified the total time spent in sleep bouts lasting more than 25 minutes, in line with what has been reported for identifying a deeper sleep stage [43]. Glial knockdown of *ruby* significantly increased long sleep bouts compared to controls (**Fig. 3D**). The increased sleep was due to longer bout lengths, as the total bout number was decreased in *ruby* knockdown flies (**Figure S3A-B**). These changes in sleep were reflected in state transition analysis when performed using a Hidden Markov Model [44], showing *ruby* knockdown significantly decreased the probability of waking (P(Wake)) and increased the probability of dozing (P(Doze)) compared to controls (**Figure S3C-D**). These findings are consistent with loss of *ruby* leading to excessive sleepiness that results in longer total sleep and prolonged sleep bouts.

**Figure 3.**
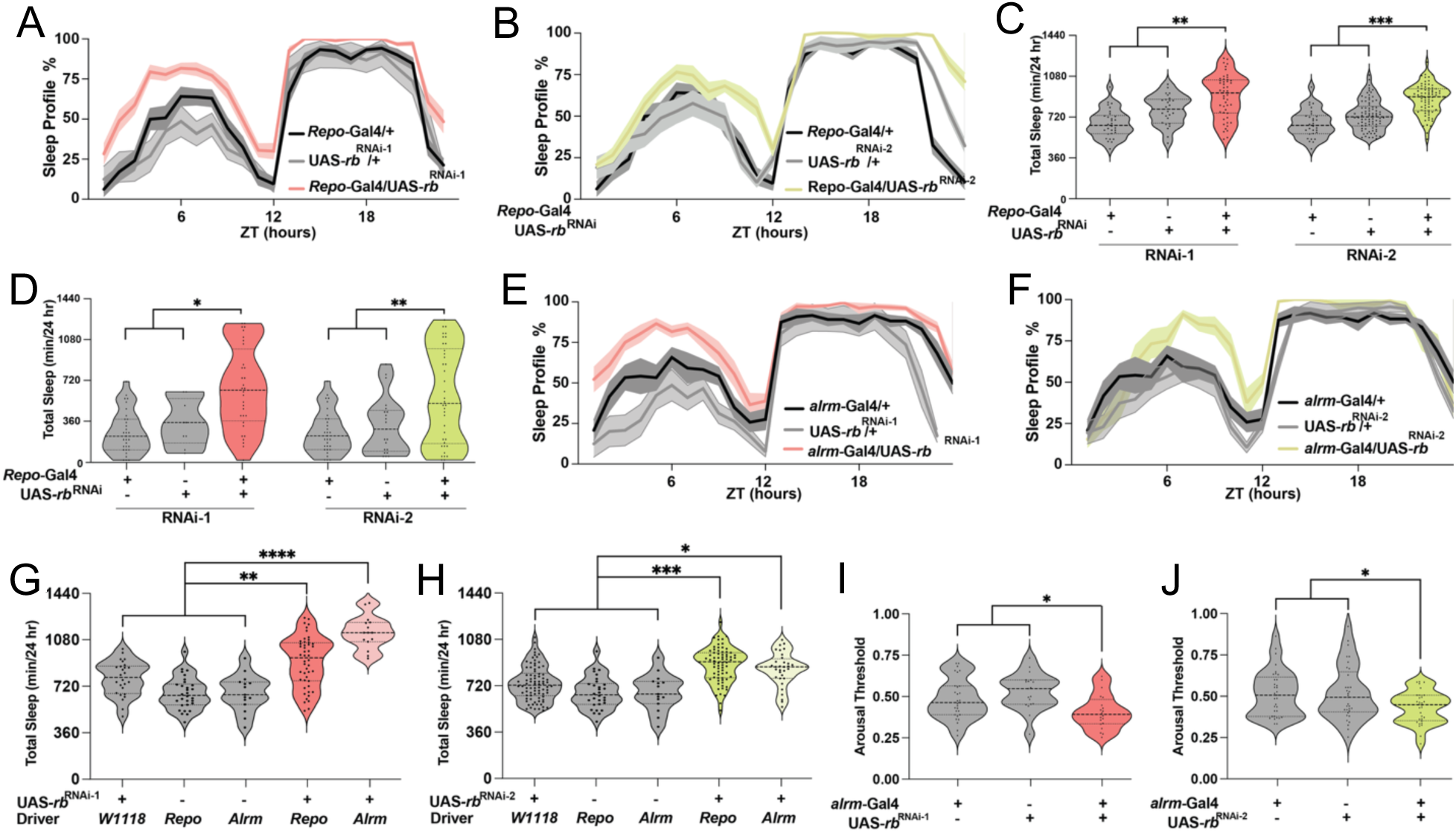
Glial knockdown of *ruby* (*ap3b2*) increases sleep. **A-B.** Sleep profile across 24 hours in flies expressing RNAi in glia under the control of *repo*-GAL4 displays an increase in sleep compared to both GAL4 and UAS controls. **C.** Total sleep (minutes over 24 hours) in two independent RNAi lines targeting *ruby* (RNAi^1^ and RNAi^2^) expressed in glia is significantly increased relative to controls (One-way ANOVA: F_2,101_ = 20.36, F_2,181_ = 37.01, p < 0.0001). **D.** Quantification of deep sleep measured as sleep bouts ≥ 25 minutes or longer in *ruby* knockdown flies (One-way ANOVA: F_2, 69_ = 15.35, F_2, 87_ = 9.635, p < 0.0001). **E-F.** Sleep profile in flies expressing *ruby* RNAi selectively in astrocyte-like glia using *alrm*-GAL4, determines an increase in sleep compared to controls. **G-H.** Knockdown of ruby in astrocyte-like glia increased total sleep compared to controls harboring the GAL4 or RNAi line alone (One-way ANOVA: F_9, 196_ = 19.28, F_5, 181_ = 6.721, p < 0.0001). **I-J.** Arousal threshold measured using DART in astrocyte-like glia knockdown of *ruby* shows a reduced arousal threshold in *ruby* knockdown flies relative to controls (One-way ANOVA: F_2,65_= 5.850, F_2,95_= 4.065, p < 0.01). Lines and error bars represent mean ± SEM; * p < 0.05, ** p < 0.01, *** p < 0.001. ZT, Zeitgeber Time (ZT0 = lights on, ZT12 = lights off).

Multiple classes of glia, including ensheathing glia, cortex glia, subperineural glia, and astrocyte-like glia have been shown to contribute to sleep regulation [31,32,34]. Using *Drosophila* glial-specific GAL4-RNAi, we sought to localize the subtype of glia where *ruby* functions to regulate sleep. Glial specific knockdown of both RNAi lines targeting *ruby* in astrocyte-like glia (alrm-GAL4>*ruby*^RNAi^) increased sleep compared to controls harboring the RNAi or driver line alone (**Fig. 3E-H and Figure S3E-F**). There was no difference between selective knockdown in astrocyte-like glia compared to pan-glia knockdown (repo-GAL4>*ruby*^RNAi)^) (**Fig. 3G-H**), suggesting the primary effect operates through astrocyte-like glia. The increased sleep was also associated with a reduced arousal threshold, as *ruby* knockdown in astrocyte-like glia responded to weaker stimuli in the *Drosophila* Arousal Tracking (DART) assay compared to controls (**Fig. 3I-J**), suggesting the longer sleep bout length may actually compensate for lighter sleep and a lack of restorative effects of sleep. Together, these findings provide evidence that *ruby/AP3B2* functions primarily in astrocyte-like glia to regulate sleep, supporting a novel role for this gene.

### Loss of *ap3b2* in zebrafish results in excessive daytime sleep

We next asked whether the ortholog to *ruby*, *AP3B2,* was conserved in function in a vertebrate model. We leveraged zebrafish, a leading model for investigating sleep [28,45,46]. Syntenic conservation of key regulatory elements is observed even in simple vertebrates, such as zebrafish [47]. This is the case for the GWAS locus of interest, where *cpeb1a* and *ap3b2* are syntenically conserved suggesting they share a regulatory interaction. Indeed, Hi-C data derived from zebrafish brain [48] indicate a distal chromatin contact between *cpeb1a* and *ap3b2* similar to the human locus (**Fig. 4A-B**) despite a large genomic distance of over 340 kilobases between gene promoters (**Fig. 4B**). Thus, we hypothesized that both genes contribute to the sleep phenotype through a shared mechanism. Variants at the locus that implicated *AP3B2* are located within *CPEB1* and it was previously assumed *CPEB1* was the causal gene at this EDS locus [3]. To test the role of both genes in a vertebrate where we observed a shared chromatin interaction, we used CRISPR-Cas9 editing in founder embryos to target the orthologs to *CPEB1* and *AP3B2*.

**Figure 4.**
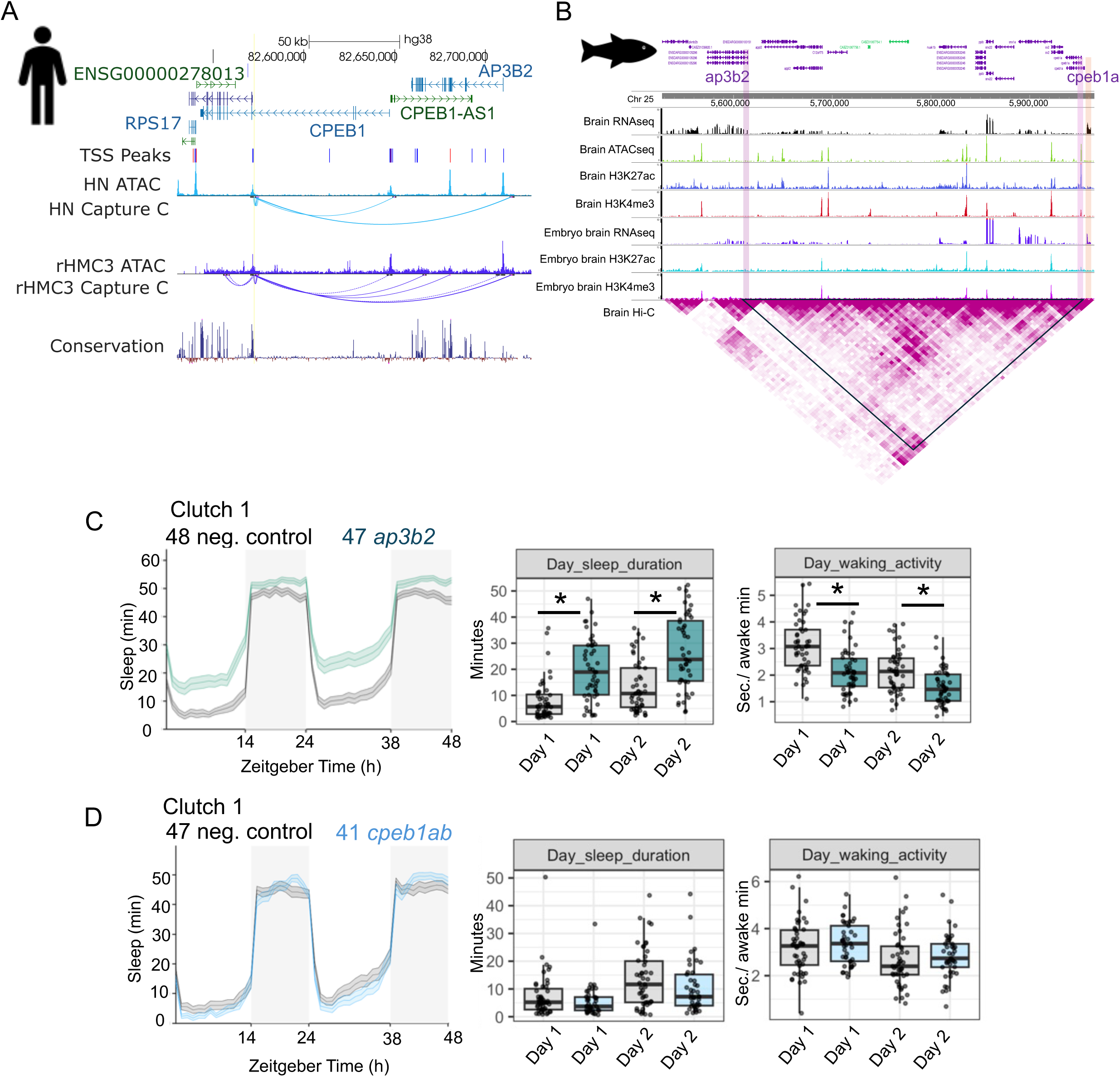
*ap3b2* is a conserved regulator of daytime sleep in zebrafish. **A.** UCSC browser plot against human hg38 genome showing the chromatin interactions between the GWAS-implicated variant (yellow line) residing in CPEB1 and its different chromatin contacts (capture C) in hypothalamic neurons (HNs) and resting microglia (rHMC3) cell lines. **B.** WashU epigenome browser plot showing the syntenic conservation between *ap3b2* and *cpeb1a* on chromosome 25 against zebrafish *danrer10* genome. Histone marks of active promoters (H3K4me3 and enhancers H3K27ac) along with chromatin accessibility (ATAC), RNA sequencing reads and chromatin interactions (Hi-C) from zebrafish brain. Purple highlight bars show promoters of *cpeb1a* and *ap3b2* and orange highlight shows unannotated region nearby *cpeb1a* contacting *ap3b2* whose sequence maps to the human *AP3B2* 3’ UTR. **C.** Sleep duration traces from *ap3b2* crispants and sibling-matched negative control-injected larvae from clutch 1. Quantification of day sleep duration and day waking activity are shown for both days of recording (6 and 7 days post-fertilization). **D.** Sleep duration traces from *cpeb1ab* crispants and sibling-matched negative control-injected larvae from clutch 1. Quantification of day sleep duration and day waking activity are shown for both days of recording (6 and 7 days post-fertilization). Box and whisker plots indicate median and quartiles. Asterisks indicate FDR-corrected significance p < 0.05 following multiple comparisons correction for 22 measures.

We designed a high-specificity single guide RNA targeting a conserved functional domain for each ortholog (**Figure S4**). Single cell embryos were injected with preformed ribonucleoprotein complexes containing the tracr:gRNA duplex and Cas9, which facilitates rapid genome editing that minimizes off-targets [49–52]. These guide RNAs proved to produce highly efficient mutations with on-target mutation rates consistently greater than 90% as measured by PCR and gel electrophoresis (**Figure S4**). Sleep and wake behaviors were measured in founder larvae using a well-established automated video-tracking program on days 6 and 7 post fertilization across two independent clutches of zebrafish larvae. *ap3b2* crispants displayed significantly increased sleep during the day in both clutches (**Figure 4C and Figure S5**) mainly driven by an increase in the number of sleep bouts (**Figure S5**). Although fish appeared free from gross morphological changes and did not show markedly reduced survival rates (**Figure S5**), their daily waking activity was significantly reduced (**Figure 4C**), indicating hypoactivity during waking episodes. Because *AP3B2* has been associated with increased neuronal activity and epileptic seizures [53], we examined sleep and activity across individual one-minute bins to identify signs of epileptic activity (e.g. rapid swimming bouts, hypermovement) [54], and we did not observe indications of these behaviors using a sensitive detection method of pixel displacement (**Figure S5E**). However, it is possible the excessive sleep is a consequence of heightened neuronal activity, which has been observed in other zebrafish models [55].

We next performed CRISPR mutation of *cpeb1a* in zebrafish and found it did not significantly influence sleep (data not shown). However, *cpeb1* is duplicated in zebrafish, so we targeted both ohnologs simultaneously to mitigate potential compensation, and again, no significant effect was observed (**Fig. 4 and Figure S6**). Together, these results support a role for *AP3B2* in sleep regulation and demonstrate the gene implicated via nearest gene mapping to the GWAS association is likely not the primary causal gene.

## Discussion

GWAS have provided an initial resource for causal gene identification; however, a major hurdle is in correct assignment of underlying causal variants to their corresponding effector genes in the appropriate cellular context. We performed cell-specific chromatin-based variant-to-gene mapping for a pathological subtype of excessive sleepiness driven by sleep fragmentation. This approach, importantly, provides cellular context for putative effector genes implicated at GWAS loci for this sleep abnormality.

Our data demonstrate there is predictive capacity in variant-to-gene mapping whereby cell-specific chromatin interactions implicate genes in the cellular context in which they are most likely to influence the trait. Specifically, we found that genes exert their influence on sleep differently depending on the cellular context. For example, *TyrRs* (*YARS1*) had opposing phenotypes when knocked down in neurons compared to glia. While some genes, such as *ruby* (*AP3B2*) only significantly influenced sleep in a single cell type. The cell-specific knockdown approach also identified potential novel cellular roles for well-known regulators of sleep. It was particularly notable that loss of *homer* in glia produced a significant sleep phenotype, while it had no effect when knocked down in neurons using *nSyb*-GAL4. Though the variant-to-gene mapping implicated *HOMER2* on chromosome 15, *HOMER1* is well-known for its role in neuronal synaptic plasticity as it relates to sleep [56–58]. However, this effect is largely driven by alternative splice isoforms that are not present in *Drosophila* [59,60]. Neuronal knockdown of *homer* using elav-GAL4, which acts in early development [61], has been shown to reduce sleep duration in flies [62], suggesting that, while this gene only has a single ortholog in *Drosophila*, it serves unique roles in different brain cells. Similar to *homer*, *ruby* (*AP3B2*), is also known for its neuronal role [63,64]. *AP3B2* plays a role in the formation of clathrin-coated synaptic vesicles involved in endocytosis [65], though we observed a primary effect with glial knockdown, suggesting it may be important for glial endocytosis like other AP-complex members [66]. AP3B2 was considered a neuron-specific AP-Complex subunit; however, single cell expression data from human [67], fish [68], and flies [69] indicate it is expressed in glia. These results demonstrate genes that were previously known for their roles in neuronal synaptic regulation, also serve important functions in glia.

While we elected to focus our further analysis on *ruby (AP3B2),* our validation screen identified numerous other regulators of sleep. These included previously established sleep and circadian regulators such as neuronal regulator, *nca* (*HPCA*) [36] and metabolic regulator *idit* (*FNDC5*) [4,70,71]. In addition, screening identified previously uncharacterized genes with short or long sleep phenotypes. For example, loss of the ortholog to *ZBTB8OS*, *archease,* in neurons or glia increases sleep. Loss of *archease* is associated with dysregulation of tRNA splicing and deficits in rapid protein synthesis [72], two cellular mechanisms that have yet to be implicated in sleep regulation. However, *archease*, mediates *XBP1* splicing, which has been shown to control sleep [73,74]. The identification of these additional genes provides candidates for further mechanistic validation in flies and other model systems.

Functional validation screens have demonstrated that some GWAS loci harbor multiple genes that contribute to a trait outcome [17,75,76]. This is likely due to the fact that genomic organization is not random and genes with shared functions can be clustered together sharing regulatory elements [77–79]. This was apparent in our *Drosophila* behavioral screen as multiple effector genes were contacted through variant-to-gene interactions at both the 1p35.1 and 15q25.2 loci, and ten of the fourteen positive hits were implicated at these two loci. Therefore, our high hit rate is likely due to variant-to-gene mapping uncovering coordinated gene programs involved in similar biological processes. For example, *archease* (*ZBTB8OS*) and *TyrRs* (*YARS1*), which were implicated at same GWAS locus, are both involved in tRNA processing and significantly altered sleep following both neural and glial knockdown.

It is also possible that variants within or near a gene may not act on the nearest gene at all. Through physical variant-to-gene mapping, we identified a variant within an intron of *CPEB1* that contacts the promoters of multiple distal genes including *AP3B2*. *CPEB1* and *AP3B2* maintain their syntenic relationship in zebrafish, and chromatin contact data indicate they may also share a regulatory interaction in zebrafish. However, through functional phenotyping of both *ap3b2* and *cpeb1* zebrafish CRISPR mutants, we determined *ap3b2* contributes an outsized effect to the EDS phenotype, while the effect of *cpeb1* was not significant. While we cannot rule out a role for *CPEB1* in human sleep regulation, our data from model organisms indicate this is not the only gene influencing sleep at the locus. In summary, our data indicate the gene contributing to the observed EDS phenotype is often a gene distal to the implicated risk variant.

More work is needed to resolve the underlying genetic mechanisms that contribute to sleep dysfunction. Here we found multiple genes associated with synaptic signaling and neuronal excitability alter sleep, including genes whose loss-of-function result in epileptic phenotypes. There is a known relationship between excessive sleepiness and seizure risk [80–82] suggesting at least some of the genetic liability to high sleep propensity may arise from the genetic predisposition to heightened neuronal activity, which may occur through neural or glial dysfunction [81,82]. Understanding genetic risk for excessive sleep is important for developing targeted therapies. It is, however, critical that genetic findings from GWAS are validated and interpreted correctly. While we cannot rule out that some implicated genes impact sleep, our findings implicate multiple novel genes in EDS. In many cases the gene identified was not the nearest gene implicated by the given GWAS locus. Therefore, GWAS results should be interpreted with caution when relying on nearest-gene assumptions.

For the first time, we demonstrate that cell-specific variant-to-gene mapping improves *in vivo* cell-type inference for genes implicated at human GWAS loci for sleep traits. Together, these studies establish a generalizable discovery framework in which GWAS-informed variant-to-gene mapping, coupled with functional validation in tractable genetic models, can identify genes and pathways underlying distinct sleep phenotypes.

## Methods

### Variant-to-gene mapping

Definition of cis-Regulatory Elements (cREs): We intersected ATAC-seq open chromatin regions (OCRs) of each cell type with chromatin conformation capture data determined by Hi-C/Capture-C of the same cell type, and with promoters (−1,500/+500bp of TSS) defined by GENCODE v40.

Genetic loci included in variant-to-genes mapping: we used the 27 significant variants that were implicated by Wang et. al. 2019 [3] to influence excessive sleepiness via sleep fragmentation. Proxies for each lead variant were queried using TopLD [83] and LDlinkR tool [84] with the GRCh38 Genome assembly, 1000 Genomes phase 3 v5 variant set, European population, and LD threshold of *r^2^*≥0.8 (results in 2206 variants).

We mapped variants to genes only when the variant resided within a defined cRE at one anchor of a chromatin loop and the distal anchor of that same loop intersected a gene promoter. This approach allowed us to link non-coding variants to distal target genes, prioritizing functional variant-gene pairs over linear proximity.

### *Drosophila* husbandry

Flies were grown and maintained on standard *Drosophila* food media (Bloomington Recipe, Genesee Scientific, San Diego, California) in incubators (Powers Scientific, Warminster, Pennsylvania) at 25°C on a 12:12 LD cycle with humidity set to 55–65%. The following fly strains were obtained from the Bloomington Stock Center: *w*^1118^ (#5905), nsyb-GAL4 (#39171), while the RNAi stock was obtained from the Vienna *Drosophila* Resource Center or the Bloomington Stock Center. The stock numbers of all lines used for screeding are described in **Table S2** unless otherwise stated. Mated females aged 3-to-5 days were used for all experiments performed in this study.

### Sleep and arousal threshold measurements in *Drosophila*

Flies were acclimated to experimental conditions for at least 24 hours prior to the start of all behavioral analysis. Measurements of sleep and arousal threshold were then measured over the course of three days starting at ZT0 using the *Drosophila* Locomotor Activity Monitor (DAM) System (Trikinetics, Waltham, MA, USA), as previously described [85–87]. For each individual fly, the DAM system measures activity by counting the number of infrared beam crossings over time. These activity data were then used to calculate sleep, defined as bouts of immobility of 5 min or more, using the *Drosophila* Sleep Counting Macro [88], from which sleep traits were then extracted. Waking activity was quantified as the average number of beam crossings per waking minute, as previously described [88]. HMM was calculated from 1 min locomotor bins, as previously described [44]. P(Wake) was measured as the probability of transition from inactivity to activity, and P(Doze) as the probability of transition from activity to inactivity in the subsequent bin. These transition probabilities were used as measures of sleep depth and sleep pressure, respectively.

Arousal threshold was measured using the DART, as previously described [89]. In brief, individual female flies were loaded into plastic tubes (Trikinectics, Waltham, Massachusetts) and placed onto trays containing vibrating motors. Flies were recorded continuously using a USB webcam (QuickCam Pro 900, Logitech, Lausanne, Switzerland) with a resolution of 800 × 600 at 5 frames/s. The vibrational stimulus, video tracking parameters, and data analysis were performed using the DART interface developed in MATLAB (The MathWorks Inc., Natick, MA). To track fly movement, raw video flies were subsampled to 1 frame/s. Fly movement, or a difference in pixilation from one frame to the next, was detected by subtracting a background image from the current frame. The background image was generated as the average of 20 randomly selected frames from a given video. Fly activity was measured as movement of greater than 3 mm. Sleep was determined by the absolute location of each fly and was measured as bouts of immobility for 5 min or more. Arousal threshold was assessed using sequentially increasing vibration intensities, from 0 to 1.2 g, in 0.3 g increments, with an interstimulus delay of 15 s, once per hour over 24 hours starting at ZT0. Measurements of arousal threshold are reported as the proportion of the maximum force applied to the platform, thus an arousal threshold of 0.4 is 40% of 1.2g.

Statistical analyses were performed in Prism (GraphPad Software 10). One-way ANOVA was used for comparisons between two or more genotypes. All post-hoc analyses were performed using Dunnett’s multiple comparisons test.

### Animal use

All experiments with zebrafish were conducted in accordance with the Institutional Animal Care and Use Committee guidelines under the appropriate institutional approved protocols (806646 and IAC 21-001154) at the University of Pennsylvania and the Children’s Hospital of Philadelphia. Breeding pairs consisted of wild-type AB and TL (Tupfel Long-fin) strains. Fish were housed in standard conditions with 14-hour:10-hour light:dark cycle at 28.5°C, with lights on at 9am (ZT0). and lights off at 11pm (ZT14).

### CRISPR/Cas9 mutagenesis in zebrafish

Highly specific guide RNAs (gRNAs) were designed using the online tool CRISPOR [90] (http://crispor.tefor.net/) with the reference genome set to “NCBI GRCz11” and the protospacer adjacent motif (PAM) set to “20bp-NGG-Sp Cas9, SpCas9-HF, eSpCas9 1.1.” gRNAs were prioritized by specificity score (>95%) with 0 predicted off-targets for sequences with up to 3 mismatches. The zebrafish sequence was obtained using Ensembl (https://useast.ensembl.org/) with GRCz11 as the reference genome. Sequence was aligned to the human amino acid sequence using DIOPT to identify the region with highest conservation, and each gRNA was designed targeting the most 5’ conserved regulatory region that was present in all transcripts. Exon 1 was skipped to avoid usage of potential alternative start codons [91]. gRNA sequences are provided in **Table S3**. AB/TL breeding pairs were set up overnight and embryos collected in embryonic growth media (E3 medium; 5mM NaCl, 0.17 mM KCl, 0.33 mM CaCl2, 0.33 mM MgSO4) the following morning shortly after lights-on. Pre-formed ribonuclear protein (RNP) complexes containing the gRNA:tracr and spCas9 HiFi enzyme were injected at the single-cell stage alternating between the gene group and scramble-injected sibling control group. Injections of scramble gRNA and gene-specific gRNA were performed in embryos from the same breeding event (clutch). Embryos were left unperturbed for one day before being transported to fresh E3 media in petri dishes (approximately 50 per dish).

### DNA extraction and PCR for genotyping zebrafish crispants

DNA extraction was performed per the manufacturer’s protocol (Quanta bio, Beverly, MA) immediately following completion of the sleep assay, as described previously [25]. Larvae were euthanized by rapid cooling on a mixture of ice and water between 2-4°C for a minimum of 30 minutes after complete cessation of movement was observed. Genotyping was performed on all individual fish at the conclusion of each sleep assay. Restriction digest PCR was used to validate mutations and confirm editing efficiency [25,52,92]. gRNAs were designed to disrupt a restriction enzyme site such that effective mutation could be detected by a lack of restriction enzyme cutting. Mutation efficiency was calculated as the ratio of the uncut:cut amplicon. Phenotype inclusion criteria was >90% mutation efficiency [25,52,92]. Following PCR confirmation, select samples were Sanger sequenced to confirm the on-target Cas9 cut 3-4 bases upstream of the Protospacer Adjacent Motif (PAM) and determine the consequence of the mutation. Primers for genotyping and restriction enzymes are listed in **Table S3**. gRNA target region flanking primers were run on a 2% agarose gel and sequence verified using Sanger sequencing prior to the assay to verify the target region was absent of polymorphisms in our AB/TL breeding pairs that may hinder genomic editing.

### Data collection and analysis for sleep phenotyping in zebrafish

All embryos and larvae were housed in an incubator at 28.5°C, with lights on at 9am (ZT0) and lights off at 11pm (ZT14) prior to data collection. Dead embryos and chorion debris were removed daily until day 5 post fertilization. On day 5, CRISPR mutants and scramble-injected sibling controls were screened for gross morphological deficits and healthy larvae were pipetted into individual wells of a 96-well plate and placed into a Zebrabox (ViewPoint Life Sciences) for automated video monitoring. Genotypes were placed into alternating rows to minimize location bias within the plate. All animals were allowed to acclimate to the Zebrabox for approximately 18 hours before beginning continuous data collection for 48 hours starting at lights on (9am) on day 6 post fertilization. Two biological replicates were run using different clutches (sibling-matched within clutch) of embryos and well placement was flipped for each experiment to minimize location bias across experiments. Each Zebrabox is sound-attenuating and contains circulating water held at a temperature of 28.5°C with automated lights cycling on the same 14-hour:10-hour light/dark cycle. Activity data were captured using automated video tracking (Viewpoint Life Sciences) software in quantization mode. As described previously [25,93], threshold for detection was set as the following: detection threshold: 20; burst: 29; freeze: 3; bin size: 60 seconds. Data were processed using custom MATLAB scripts [94] to calculate the following parameters for both day and night separately: sleep duration (minutes/hour), activity duration (seconds/hour), waking activity (seconds/awake minute/hour), sleep bout length (minutes/bout), sleep bout number (number/hour) and nighttime sleep latency (minutes). Statistical analysis was performed in R. Data are presented for both day/night 6 and 7 post fertilization. Strictly standardized mean difference[95] was calculated across sleep traits for each crispant group relative to its sibling-matched controls. Clutches were analyzed separately. Significance was determined by two methods, multiple unpaired t-tests or Wilcoxon rank-sum (to address non-normality) for each comparison followed by FDR correction using the Benjamini-Hochberg method. SSMD plots are shown with asterisks indicating measures that were significant (p < 0.05) by correction of the Wilcoxon rank-sum value given many sleep measures are non-parametric and this test is more robust to outliers. Statistical comparisons from both tests are largely consistent given the large sample sizes and both are provided in the **Associated Data File**.

## Web Resources

DIOPT https://www.flyrnai.org/cgi-bin/DRSC_orthologs.pl

ensembl https://useast.ensembl.org/

UCSC genome browser https://genome.ucsc.edu/

WashU epigenome browser https://epigenomegateway.wustl.edu/

## Acknowledgements

The work was supported by NIH grant T32 HL170968, R01 HL178074, P01 HL094307, and P01 HL160471. S.F.G is supported by NIH awards R01 AG057516 and R01 HD056465 and the Daniel B. Burke Endowed Chair. Authors would like to thank the CHOP Zebrafish Core for assistance in zebrafish care, maintenance, and CRISPR experimentation.

## Declaration of Interests

The authors declare no competing interests.

## Notes

### Competing Interest Statement

The authors have declared no competing interest.

