## Supplementary Figures for "Cell-specific variant-to-gene mapping identifies conserved neural and glial regulators of sleep"

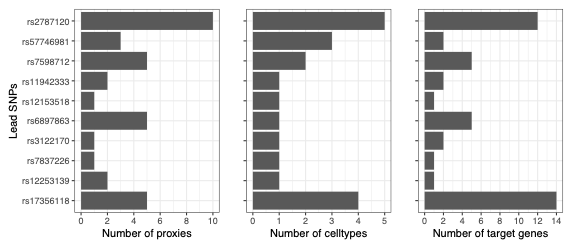

**Figure S1.**

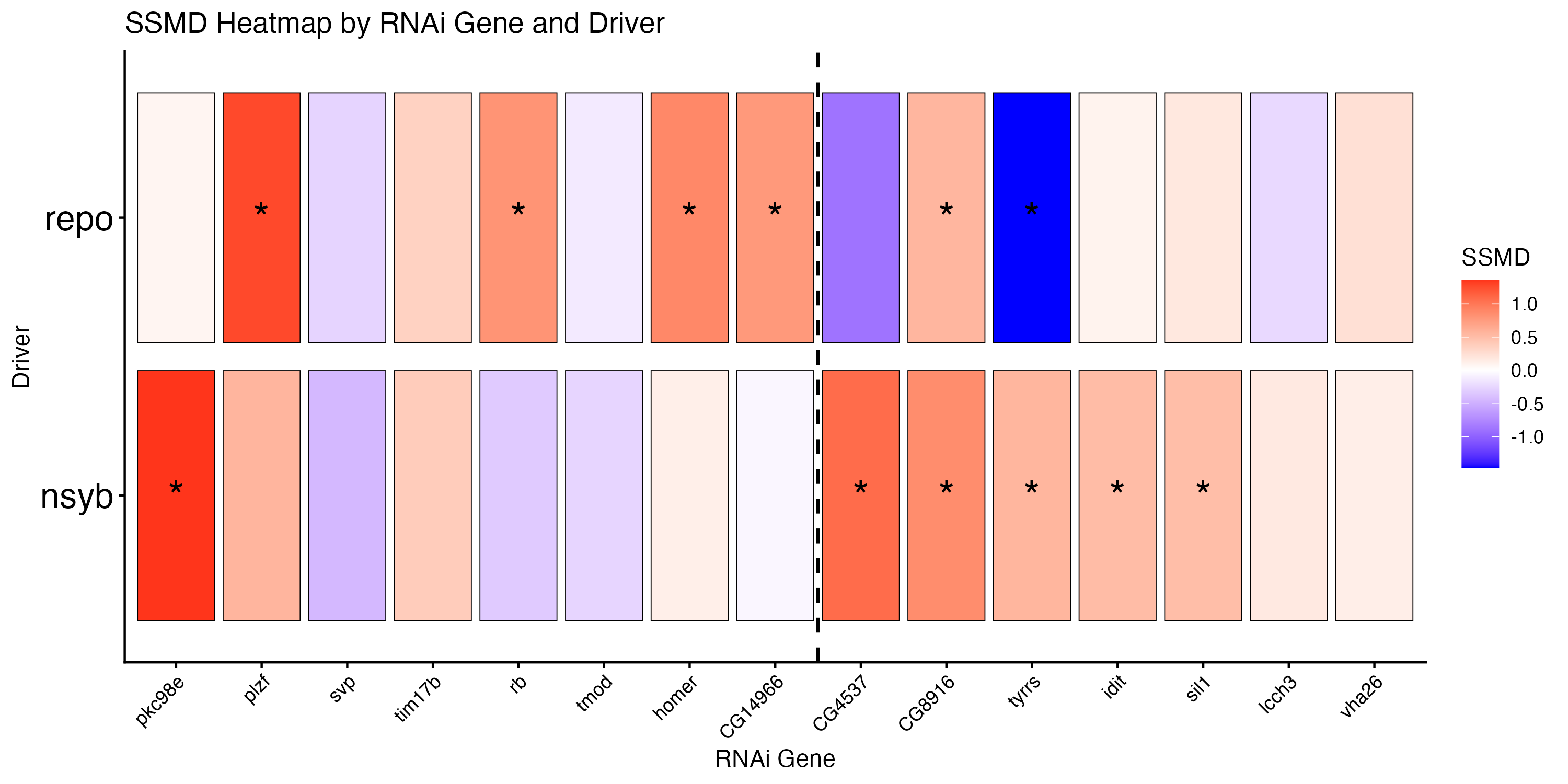

Glial genes

Neural genes

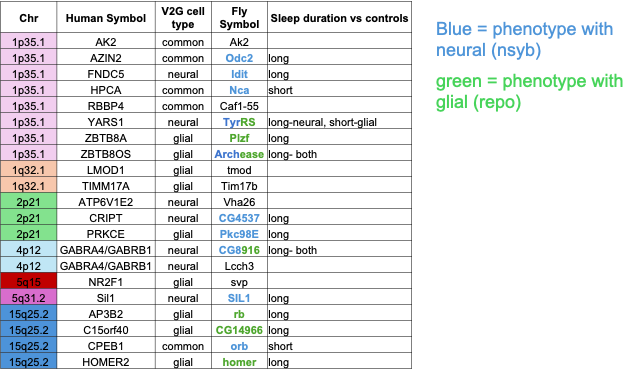

**B**

**A**

**Figure S2.**

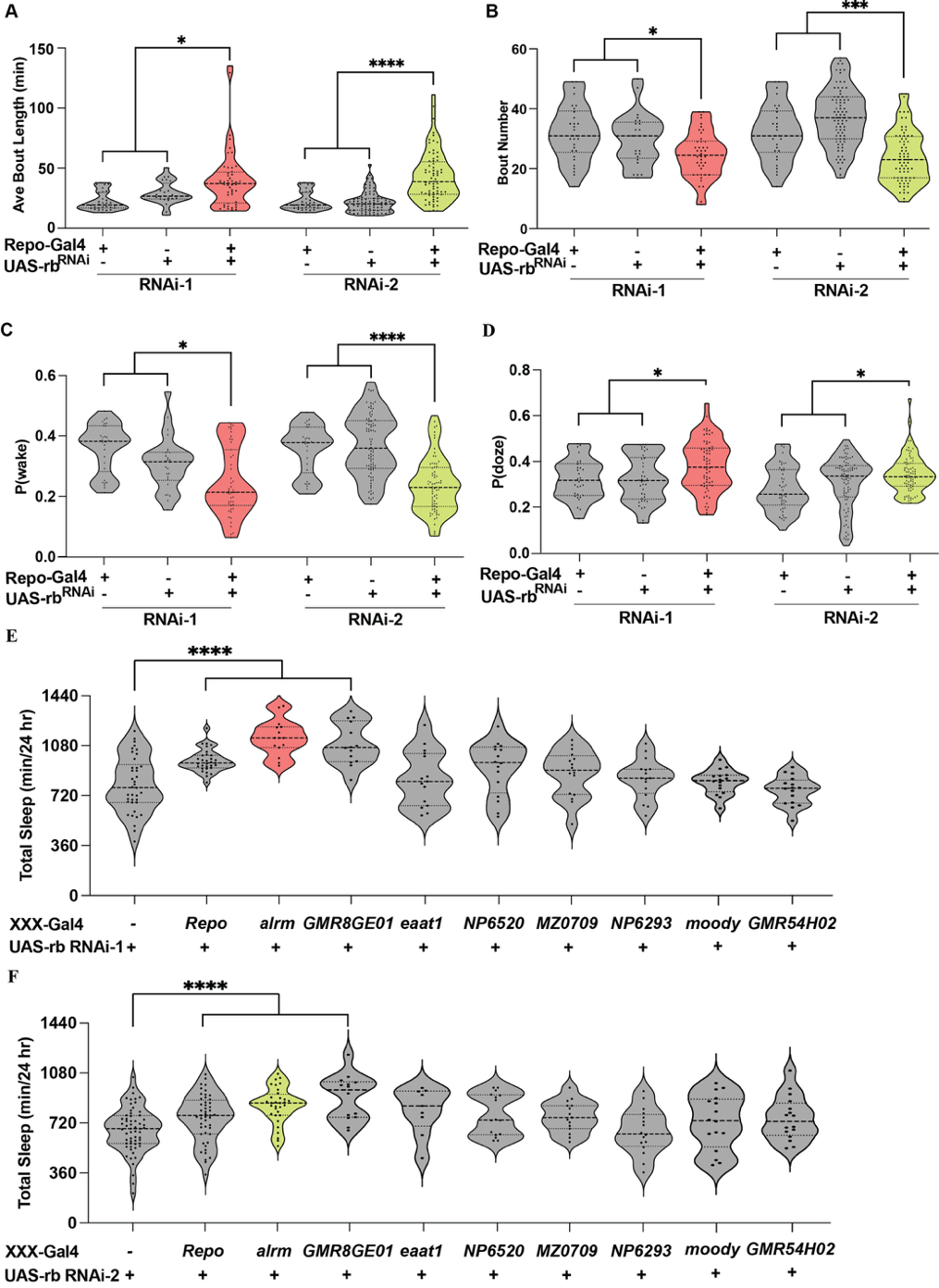

**B**

**A**

**D**

**C**

**E**

**F**

**Figure S3.**

**F**

**E**

**D**

**C**

**B**

**A**

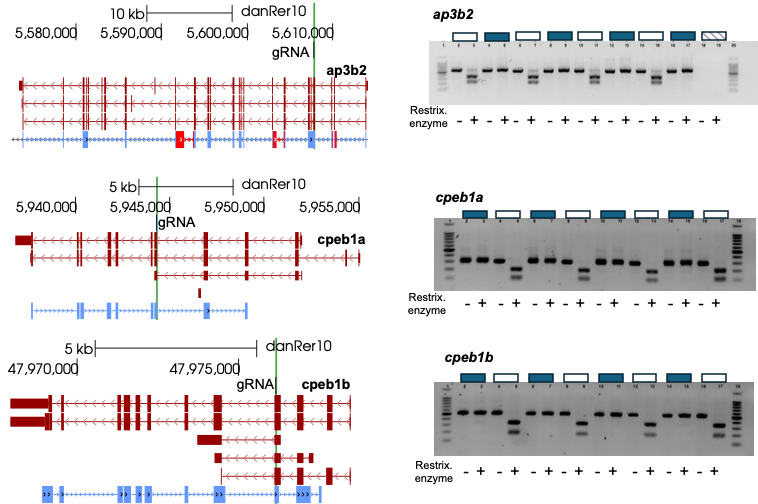

**Figure S4.**

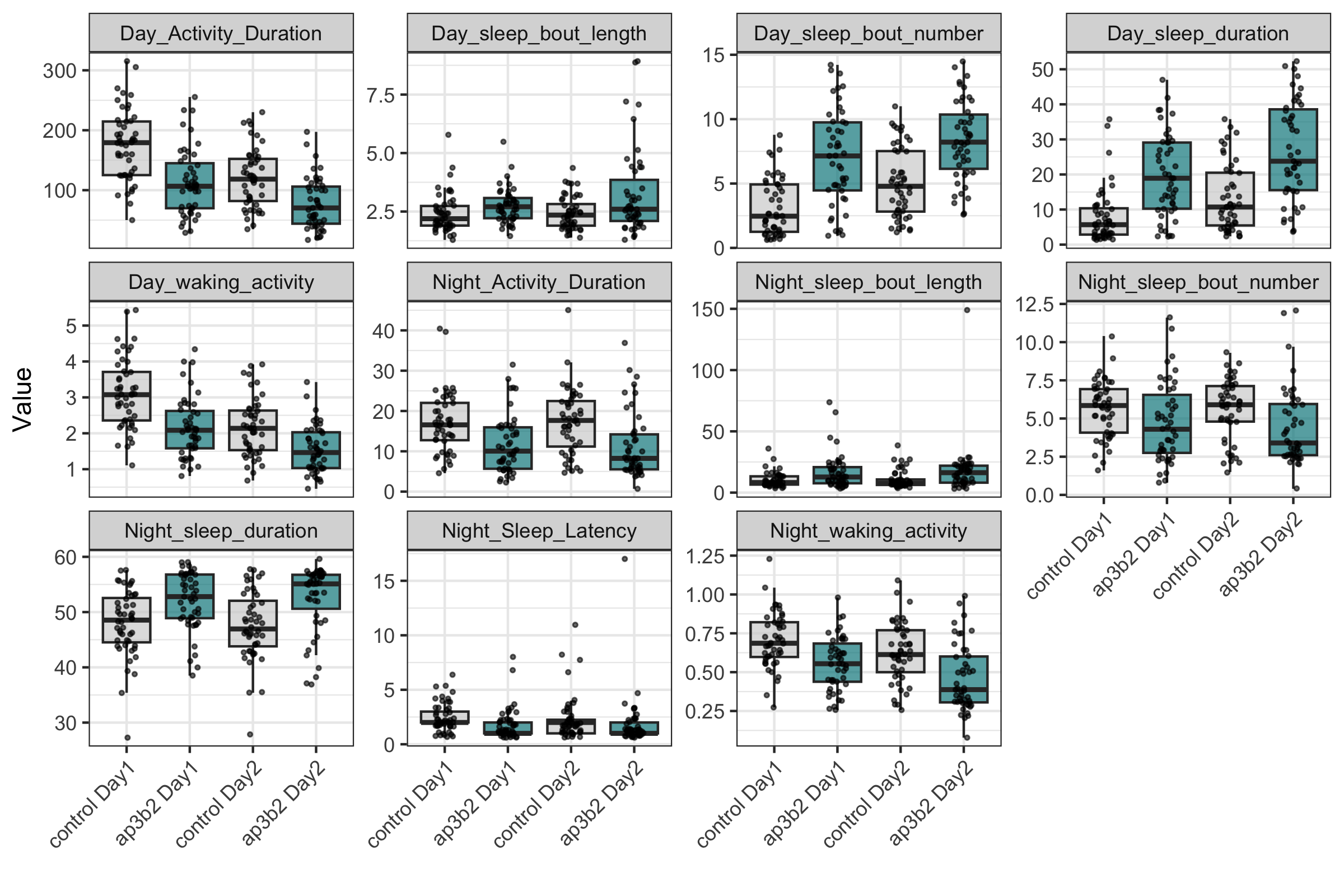

**A**

**B**

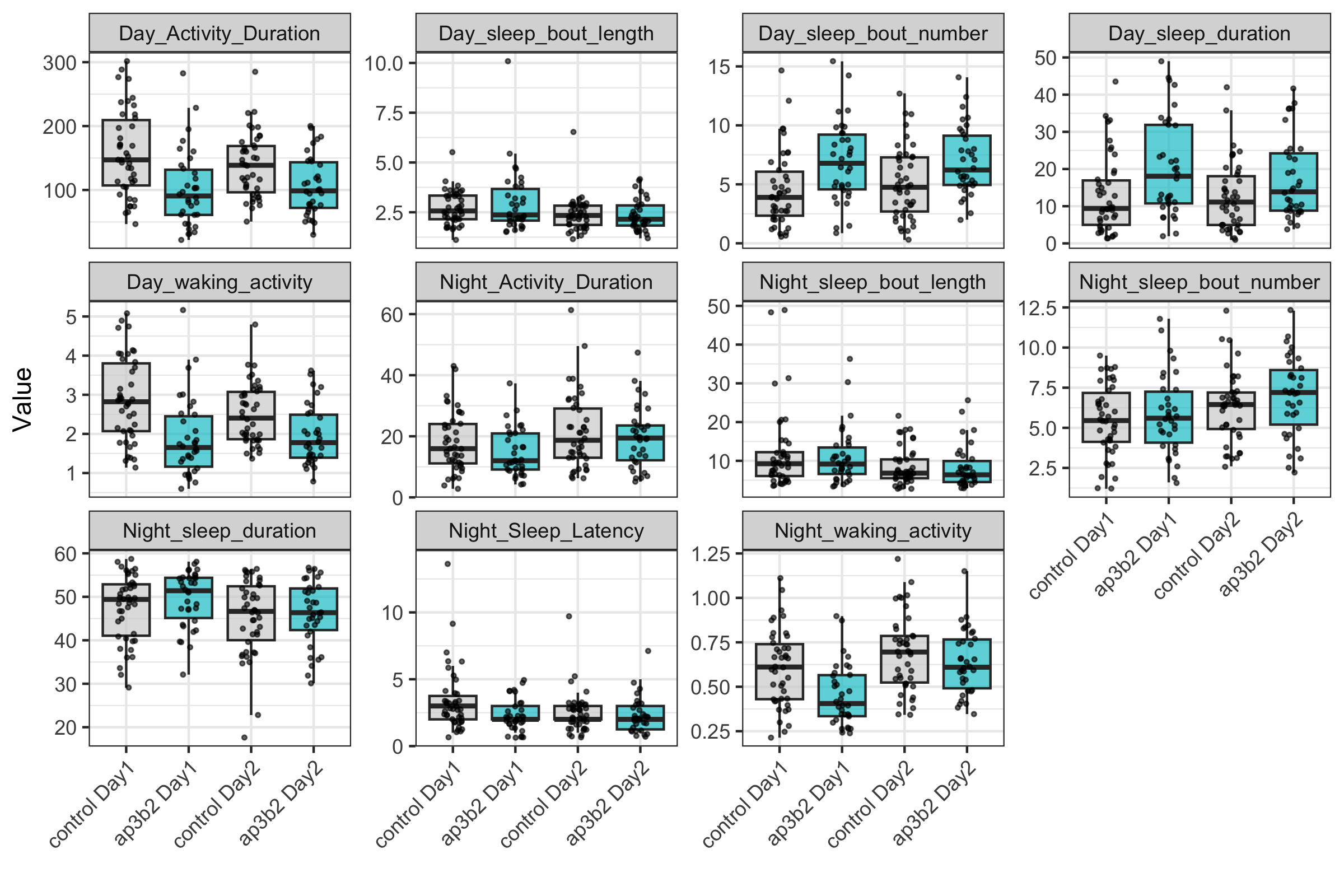

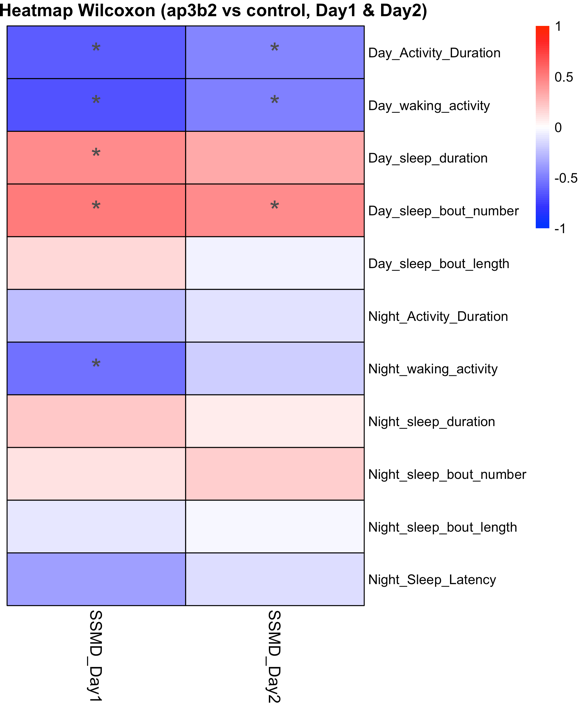

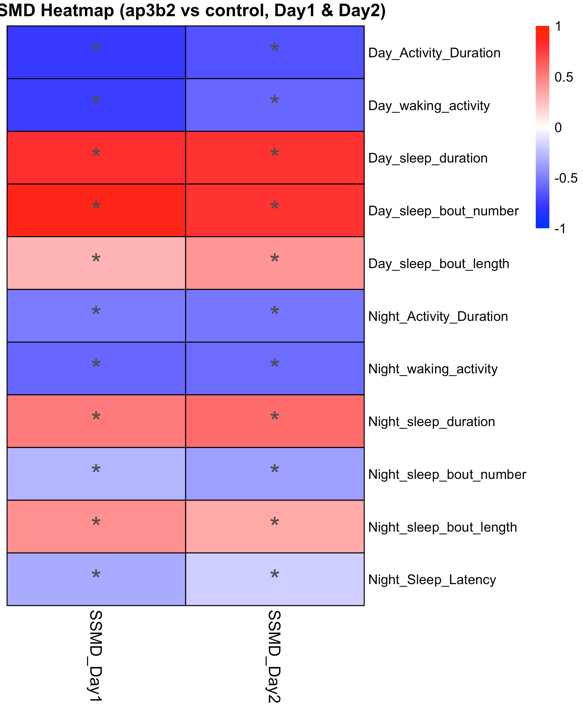

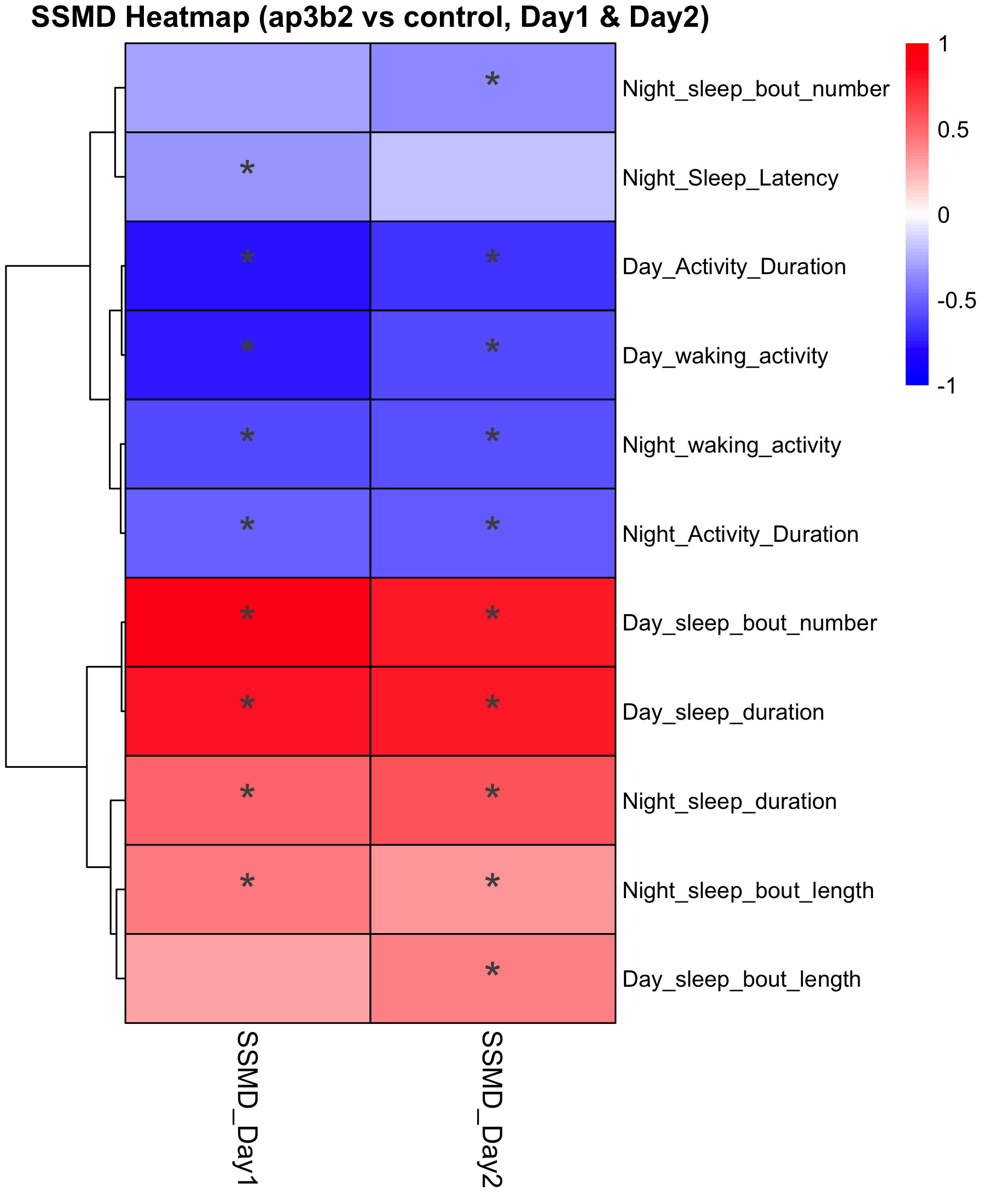

**E**

**D**

**C**

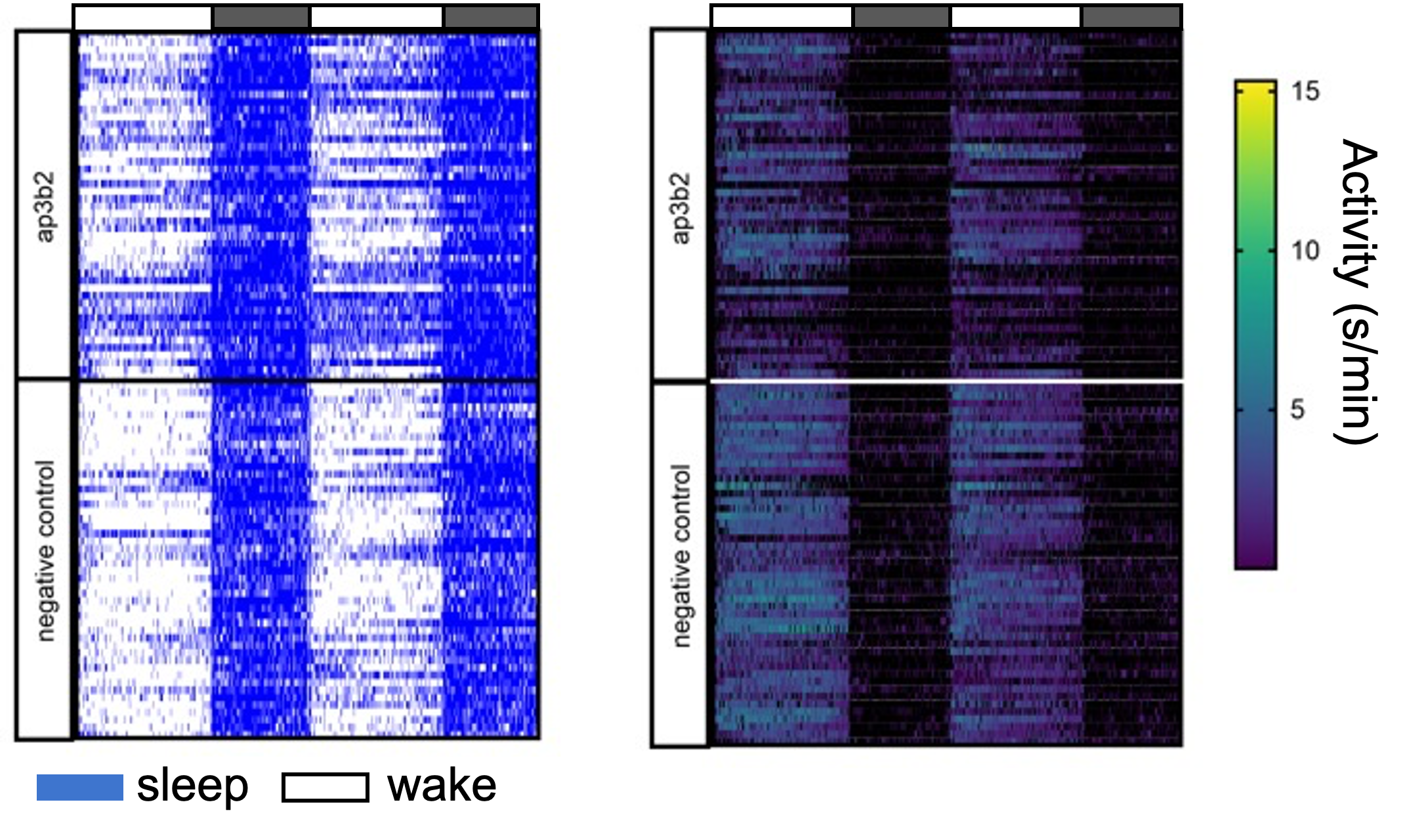

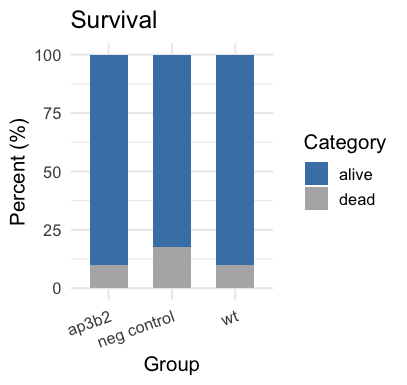

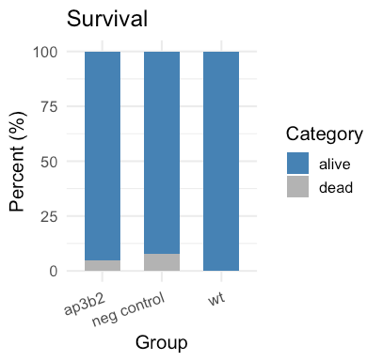

**Figure S5.**

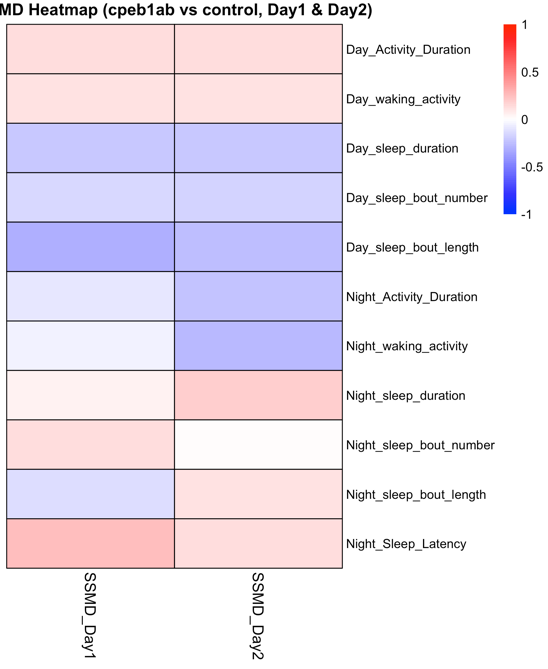

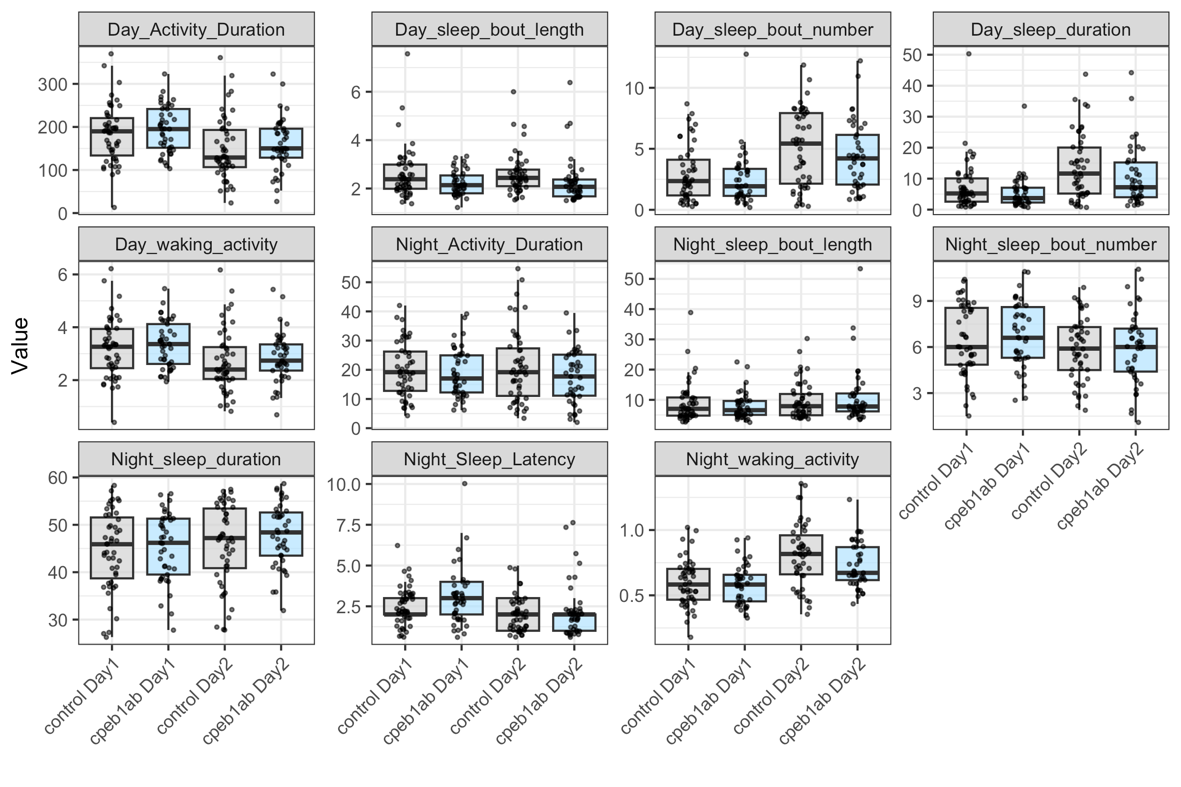

**A**

**B**

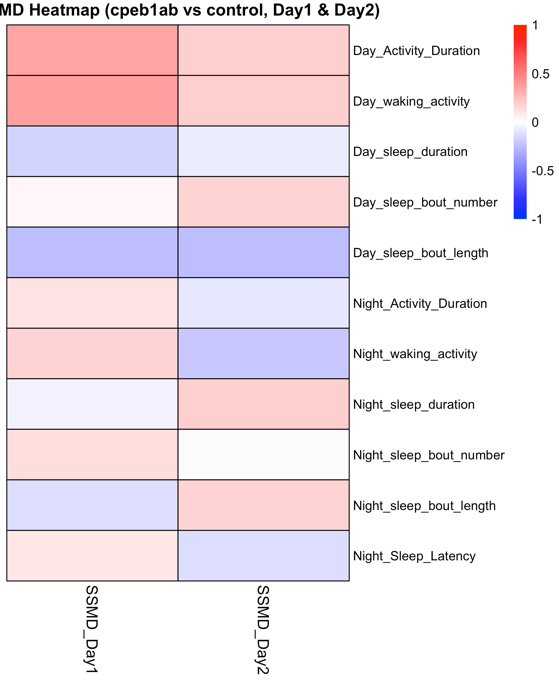

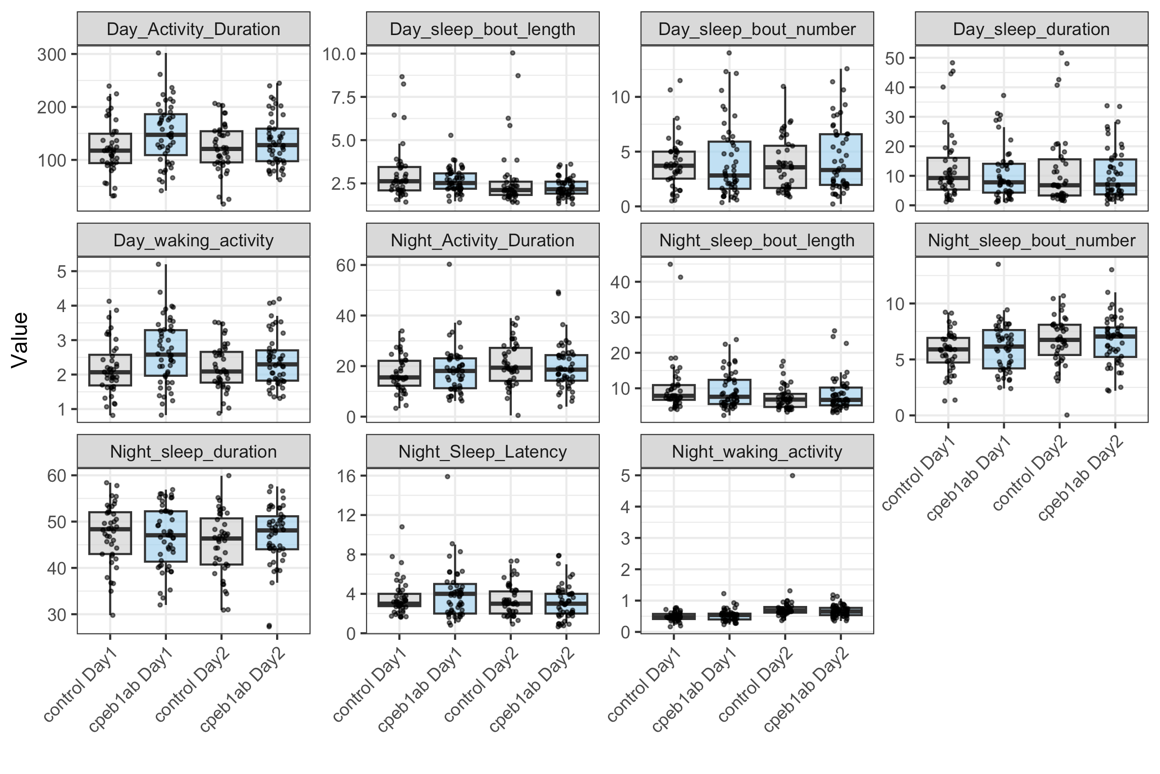

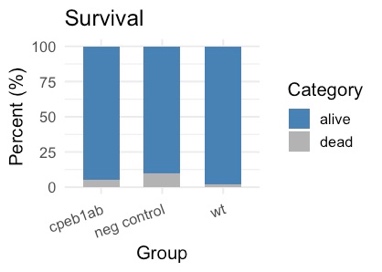

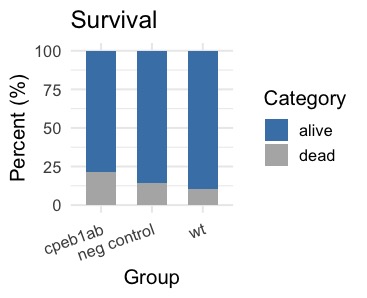

**D**

**C**

**Figure S6.**

**Supplementary Figure Legends**

**Figure S1. Variant-to-gene mapping using EDS lead SNPs.** Ten lead SNPs were obtained from Wang et al. 2019. Proxy variants (*r^2^* ≥ 0.8) that resided in defined cREs (defined in methods) were identified for each of five cell types. Target effector genes were identified in each cell type provided their promoter was accessible by ATACseq and contacted by a cRE.

**Figure S2.** **A.** Heatmap showing strictly standardized mean difference (effect size) for total sleep duration of each RNAi relative to pooled controls. Glial/neural genes indicated by the top bar refer to which cell type the gene was implicated in using variant-to-gene mapping. The GAL4 driver is shown to the left. nsyb = pan-neuronal, repo = pan-glial. Asterisk indicates significance as determined by one-way ANOVA relative to controls followed by Dunnett’s multiple comparisons test as indicated in Figure 2. **B.** Table outlining the chromosome (chr) locus, effector gene implicated shown as the human gene symbol, cell type implicating the gene from variant to gene mapping, fly ortholog from DIOPT, and effect on sleep duration relative to controls. For variant-to-gene (V2G) cell type, common refers to both a neural and a glial cell type, neural refers to human stem-cell-derived hypothalamic neural progenitors or mature hypothalamic neurons, glial refers to HMC3 microglia (activated or resting) and astrocytes. Under Fly Symbol, green indicates a significant phenotype was observed with *repo*-GAL4, blue indicates a significant phenotype was observed with *nSyb*-GAL4. Blue and green indicates a significant phenotype was observed with both *repo* and *nSyb*.

**Figure S3.** **Glial knockdown of *ruby* alters sleep. A.** Average bout length in *ruby* glial knockdown using *repo*-GAL4 determines a significant increase compared to controls (F_2,99_ = 9.273, F_2, 181_ = 46.69, p < 0.0001). **B.** Sleep bout number in *ruby* knockdown flies show a reduction relative to controls (F_2, 94_ = 6.904, F_2, 181_ = 38.86, p < 0.0001). **C-D.** Hidden Markov Model (HMM) analysis of sleep state reveals that the probability of waking (P(Wake)) is significantly decreased in ruby RNAi lines compared to controls (F_2, 100_ = 12.18, F_2, 181_ = 40.55, p < 0.0001). The probability of dozing (P(Doze)) is significantly increased in *ruby* RNAi lines (F_2, 128_ = 5.276, F_2, 204_ = 6.457, p < 0.001). **E-F.** Knockdown of *ruby* in glial subpopulations revealed a significant increase in total sleep in astrocyte-like (alrm and GMR86E01) glial drivers. These GAL4 drivers were used: eaat1-GAL4, NP6520-GAL4, and MZ0709-GAL4 (ensheathing glia), NP6293-GAL4 (perineurial glia) and GMR54H02-GAL4 (Cortex glia), moody-GAL4 (Subperineurial glia). (F _11, 226_ = 17.90, F _9, 232_ = 5.304, p < 0.0001). Lines and error bars represent mean ± SEM; * p < 0.05, ** p < 0.01, *** p < 0.001.

**Figure S4. CRISPR gRNA target site and genotype validation in crispants. A-C.** CRISPR gRNAs were designed to target highly conserved functional protein-coding domains that were present in all coding transcripts. gRNA location is shown above each track with PAM site highlighted in green for *ap3b2* (A), *cpeb1a* (B), and *cpeb1b* (C). All transcripts are shown in red with orientation indicated by arrow direction. Blue track on the bottom of each shows conservation with respect to the human genome. **D-F.** Representative blots showing PCR results run on 2% agarose gel using target-specific primers. Negative controls are shown with white bars and gRNA-injected fish are shown with blue bars. Dashed gray lines in end well of D indicate water control. Molecular ladder is shown on both sides indicating 100bp intervals. Presence of target specific restriction enzyme is indicated below each blot. In mutant fish, the restriction enzyme site is deleted resulting in no digestion as indicated by equal size bands in adjacent – and + wells. If the sequence is wild-type, the restriction enzyme will cut resulting in digestion and bands of distinct sizes in adjacent – and + wells. The ratio of the two bands is used to determine cutting efficiency. Fish were included in the assay if their mutation efficiency was >90%. *cpeb1a* and *cpeb1b* blots are showing the same 8 fish at both target locations.

**Figure S5.** **All sleep wake measures in *ap3b2* crispants.** Clutch 1 (**A**) and Clutch 2 (**B**) sleep and activity measures for both measured days (days 6-7 post fertilization). Individual animals are shown as points, and box and whisker plots indicate median and quartiles. Strictly standardized mean difference (SSMD) plots are shown at the right indicating effect size relative to sibling and camera-matched controls. Asterisks indicate FDR-corrected *p*-value p < 0.05 following Wilcoxon rank-sum test. Survival rates are shown for clutch 1 (**C**) and clutch 2 (**D**). All fish (*ap3b2*, neg. control, and wt come from the same clutch). **E.** Minute-by-minute analysis of behavior in ap3b2 crispant fish and negative controls. Each row represents a fish and each column represents a one-minute bin across 48 hours. Sleep and wake are indicated by blue and white, respectively, and activity is shown as total moving activity (seconds per minute). White bars along top indicate day and gray bars indicate night.

**Figure S6. All sleep wake measures in *cpeb1ab* crispants**. Clutch 1 (**A**) and Clutch 2 (**B**) sleep and activity measures for both measured days (days 6-7 post fertilization). Individual animals are shown as points, and box and whisker plots indicate median and quartiles. Strictly standardized mean difference (SSMD) plots are shown at the right indicating effect size relative to sibling and camera-matched controls. Asterisks indicate FDR-corrected *p*-value <0.05 following Wilcoxon rank-sum test. Survival rates are shown for clutch 1 (**C**) and clutch 2 (**D**). All fish (*cpeb1ab*, neg. control, and wt come from the same clutch).
